# RNA processing by the CRISPR-associated NYN ribonuclease

**DOI:** 10.1101/2024.04.08.588522

**Authors:** Haotian Chi, Malcolm F White

## Abstract

CRISPR-Cas systems confer adaptive immunity in prokaryotes, facilitating the recognition and destruction of invasive nucleic acids. Type III CRISPR systems comprise large, multisubunit ribonucleoprotein complexes with a catalytic Cas10 subunit. When activated by the detection of foreign RNA, Cas10 generates nucleotide signalling molecules that elicit an immune response by activating ancillary effector proteins. Among these systems, the *Bacteroides fragilis* type III CRISPR system was recently shown to produce a novel signal molecule, SAM-AMP, by conjugating ATP and SAM. SAM-AMP regulates a membrane effector of the CorA family to provide immunity. Here, we focus on NYN, a ribonuclease encoded within this system, probing its potential involvement in crRNA maturation. Structural modelling and *in vitro* ribonuclease assays reveal that NYN displays robust sequence-nonspecific, Mn^2+^-dependent ssRNA-cleavage activity. Our findings suggest a role for NYN in trimming crRNA intermediates into mature crRNAs, which is necessary for type III CRISPR antiviral defence. This study sheds light on the functional relevance of CRISPR-associated NYN proteins and highlights the complexity of CRISPR-mediated defence strategies in bacteria.

## Introduction

CRISPR-Cas (Clustered Regularly Interspaced Short Palindromic Repeats-CRISPR associated proteins) systems provide anti-viral adaptive immunity in prokaryotes, capable of memorising past invasions and targeting the same invading nucleic acids upon future infections [1, 2]. The CRISPR immune response typically involves three stages: adaptation, expression, and interference. In the adaptation stage, foreign nucleic acids are captured and subsequently integrated into the CRISPR array between two repeat sequences [3, 4]. During expression, Cas proteins are expressed and the CRISPR array is transcribed into a precursor crRNA (pre-crRNA), which is subsequently processed into mature CRISPR RNAs (crRNAs) [5]. These crRNAs, containing a partial repeat sequence and a single spacer sequence, play a vital role in guiding Cas proteins during the interference stage to detect and eventually degrade foreign nucleic acids [6].

While these general immune responses are shared among CRISPR-Cas systems, the detailed mechanisms of action vary significantly. Consequently, CRISPR-Cas systems have been classified into 2 classes, 6 types and more than 30 subtypes [7–9]. The key distinction between these two classes lies in the composition of their interference effector: class one systems (type I, III and IV) feature multiple subunits, while a single multidomain effector protein is present in class two systems (type II, V and VI). Class one systems are more prevalent, found in more than half of the CRISPR loci in both bacteria and archaea [8].

Type III CRISPR systems stand out for their reliance on signalling pathways to confer complex and multifaceted immune response [10]. Most subtype III-A (Csm, Cas subtype Mtube) and III-B (Cmr, Cas module RAMP) systems exhibit the capability to generate cyclic oligoadenylates (cOA) as second messengers, facilitated by the enzymatic subunit Cas10, which contains two polymerase-cyclase Palm domains [11, 12]. Upon detecting viral RNAs, cOAs are generated with 3 to 6 AMP monomers joined with 3’-5’ phosphodiester bonds. These second messengers subsequently activate downstream accessory proteins to non-specifically degrade RNA, DNA, or other cellular components, ultimately resulting in viral clearance, cell dormancy or death [13]. The CARF (CRISPR-Associated Rossmann Fold) domain is an extensively studied cOA sensor domain in accessory proteins, some of which also exhibit cOA degradation activity, acting as self-limiting accessory proteins or ring nucleases to regulate signalling pathways [14–18]. Interestingly, viruses have also been found to utilise ring nucleases as an efficient anti-CRISPR strategy [19, 20].

We recently characterised a type III-B CRISPR system from *Bacteroides fragilis*, which includes three non-characterised ancillary effectors: a CorA putative divalent cation channel protein, a DHH/DHHA1-family phosphodiesterase of the NrN family [21], and a <u>N</u>edd4-BP1, <u>Y</u>acP <u>N</u>uclease (NYN) family nuclease, denoted as CorA, NrN and NYN, respectively [22–24]. Our previous work has shown that upon binding to target RNA, the *B. fragilis* Cmr complex is activated to synthesise a new class of signal molecule SAM-AMP through conjugation of SAM (S-adenosyl methionine) with ATP catalysed by Cas10 subunit [23]. The accessory membrane protein CorA, activated by SAM-AMP binding, is thought to disrupt the cell membrane integrity, leading to cell dormancy or death to prevent spreading of phage infection [23]. The accessory protein NrN was shown to degrade SAM-AMP, presumably acting in an equivalent role to ring nucleases to switch-off the signalling pathway. However, the role of the third ancillary effector NYN in the *B. fragilis* CRISPR systems remains unclear.

CRISPR-associated NYN is a member of the NYN domain family of ribonucleases [22, 25]. The NYN domain belongs to the PIN domain-like superfamily, a major metal-dependent nuclease superfamily widely present in all three domains of life [25, 26]. Proteins containing the PIN-like domain participate in various cellular processes, including DNA replication and repair, mRNA decay, rRNA maturation and toxin-antitoxin systems [26]. For example, Rael (YacP) is involved in mRNA decay in a ribosome-dependent manner in *Bacillus subtilis* [27] and the N-terminal NYN domain of Marf1 (meiosis regulator and mRNA stability factor 1) exhibits ribonuclease activity, essential for oocyte meiosis and genome integrity [28]. However, no NYN family proteins have been characterised in the context of CRISPR systems.

Here, we investigate the potential functional roles of NYN in the *B. fragilis* type III-B CRISPR system and demonstrate its manganese-dependent ribonuclease activity, which non-specifically degrades a wide range of single-strand RNAs *in vitro*. The properties of *B. fragilis* NYN are consistent with a role in crRNA maturation in type III CRISPR systems.

## Results

### Investigation of NYN ribonuclease function in the *B. fragilis* Cmr system

The gene encoding NYN is situated adjacent to the genes encoding Cas6, CorA and NrN within the *B. fragilis* type III-B CRISPR locus [29] (Figure 1A). The functional correlation of NYN within the context of *B. fragilis* Cmr systems aroused our interest, given the fact that colocalised genes in prokaryotic operons often have a functional connection. Our previous findings demonstrated that the *B. fragilis* Cmr system provides immunity via SAM-AMP signalling in the heterologous host *E. coli* when both ancillary effectors CorA and NrN are present [23], suggesting an auxillary function for the NYN protein. We first investigated whether NYN serves as an alternative pathway for SAM-AMP turnover. To investigate this possibility, NYN was co-expressed in *E. coli* with the activated *B. fragilis* Cmr system to assess its impact on SAM-AMP production *in vivo*. Cells were co-transformed with plasmids pBfrCmr1-6, pBfrCRISPR_Tet and either pRATDuet-NYN or pRATDuet-NrN (as a control) to activate the system. After growth at 37 °C with full induction, cellular nucleotides were extracted and purified for HPLC analysis. The presence of NYN along with the activated Cmr system did not affect the production of SAM-AMP (Figure 1B). In contrast, SAM-AMP was absent in the presence of the phosphodiesterase NrN, which degrades SAM-AMP into SAM and AMP [23]. To investigate the activities of NYN *in vitro*, we purified *B. fragilis* NYN to homogeneity (Supplementary Figure 1). NYN protein was incubated with SAM-AMP *in vitro* in the presence of Mn^2+^. Subsequent HPLC analysis revealed no observable degradation of SAM-AMP (Figure 1C). These findings indicate that NYN is not involved in SAM-AMP metabolism.

**Figure 1.**
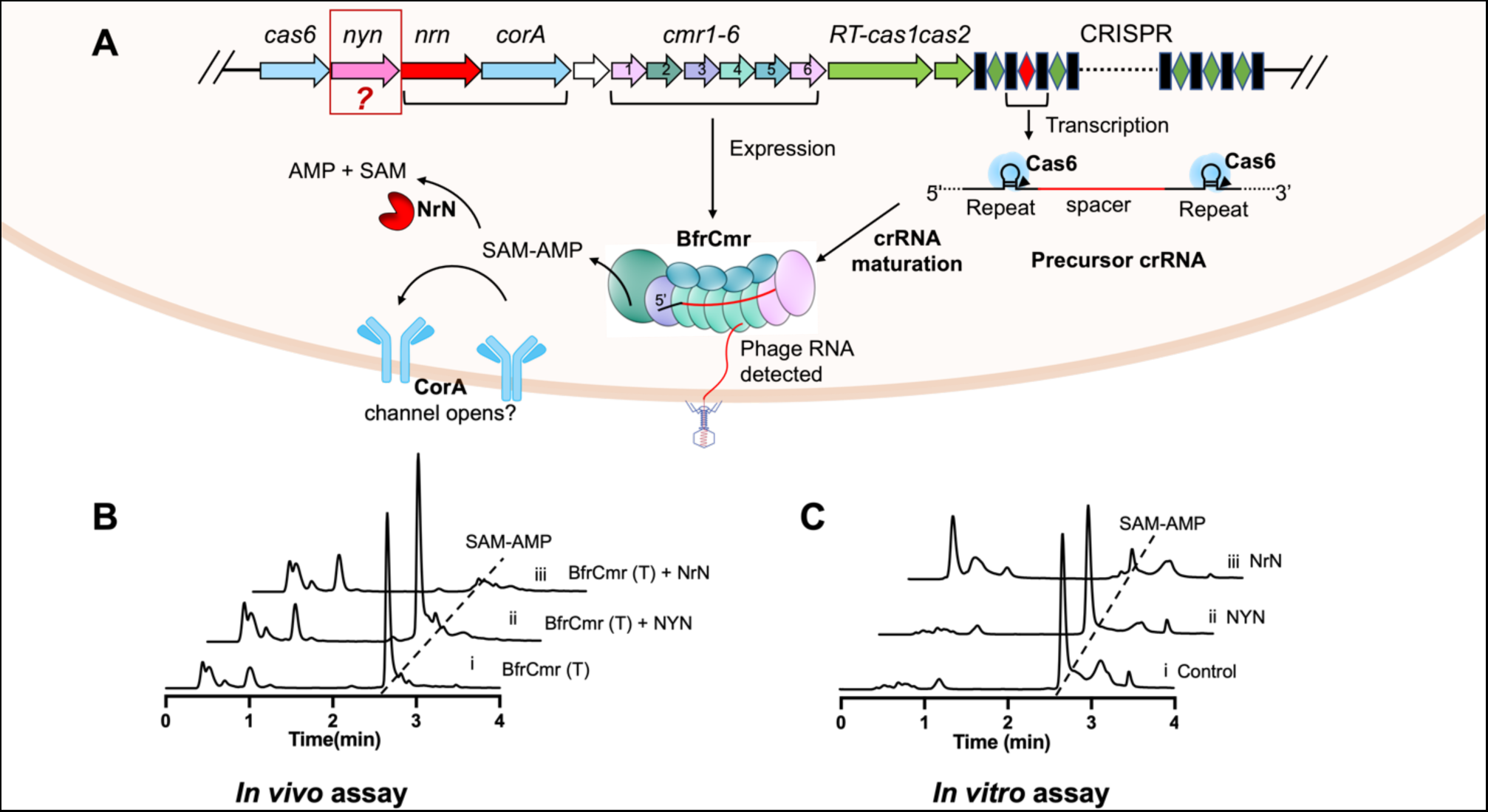
NYN investigated in the context of *B. fragilis* Cmr system. **A**. Schematic illustration of the *B. fragilis* type III CRISPR immune signalling pathway. During crRNA biogenesis, the CRISPR array is transcribed into a precursor crRNA, subsequently processed into mature crRNAs by Cas6 and other unidentified nucleases. The Cmr complex is thought to assemble around the crRNA, forming a functional Cmr-crRNA complex. Upon detection of invading RNAs during interference, the Cmr complex is activated to produce the signal molecule SAM-AMP, which in turn activates the membrane effector CorA, presumably leading to cell death by disrupting membrane integrity. NrN is involved in regulating the turnover of SAM-AMP to fine-tune the immune signalling pathway. Notably, the adjacent gene *nyn*’s function within the *B. fragilis* CRISPR immune system remains unknown. **B.** HPLC analysis of *E. coli* extracts expressing the wild-type *B. fragilis* Cmr system with target crRNA (BfrCmr (T)), along with expression of NYN or NrN. The introduction of NYN into the activated wild type BfrCmr system had no effect on the production of signal molecule SAM-AMP (trace ii), while SAM-AMP was undetectable in cells expressing NrN (trace iii). **C.** HPLC analysis of samples in which purified SAM-AMP was incubated with NYN *in vitro* (trace ii). SAM-AMP incubated with NrN served as a positive control (trace iii), while sample lacking any enzymes serves as a negative control (trace i). No degradation of SAM-AMP was observed in the presence of NYN.

### Mn^2+^-dependent ribonuclease activity of NYN

To investigate whether NYN exhibited ribonuclease activity, we incubated the purified recombinant protein with a range of RNA species and analysed reaction products by gel electrophoresis. NYN effectively cleaved the *B. fragilis* CRISPR repeat RNA (BfrCRISPR), labelled with a 5’-FAM fluorophore, resulting in the accumulation of small RNA degradation products (Figure 2A). Notably, this activity was strictly dependent on the presence of Mn^2+^, with no observed activity in the presence of Mg^2+^. This contrasts with the Marf1 NYN domain protein, which is active with either Mn^2+^ or Mg^2+^ [28, 30]. We proceeded to test the activity of NYN against a range of 5’-FAM labelled RNA oligonucleotides of different sequence and length (17-60 nt) (Figure 2B, Supplementary Figure 2). Cleavage products were distributed in the substrates and did not appear to interconvert, consistent with endonuclease activity and with previous studies of Marf1 [28, 30]. Mapping of the cleavage sites did not reveal a consensus sequence motif, suggesting relaxed sequence specificity (Supplementary Figure 2). To investigate this further, we incubated NYN with a polyuridine RNA oligonucleotide with a 3’-FAM label (PolyU_10_-FAM) (Figure 2C). This confirmed endonuclease activity for NYN, with non-interconverting cleavage products centred 3-4 nucleotides from the 3’ end of the substrate.

**Figure 2.**
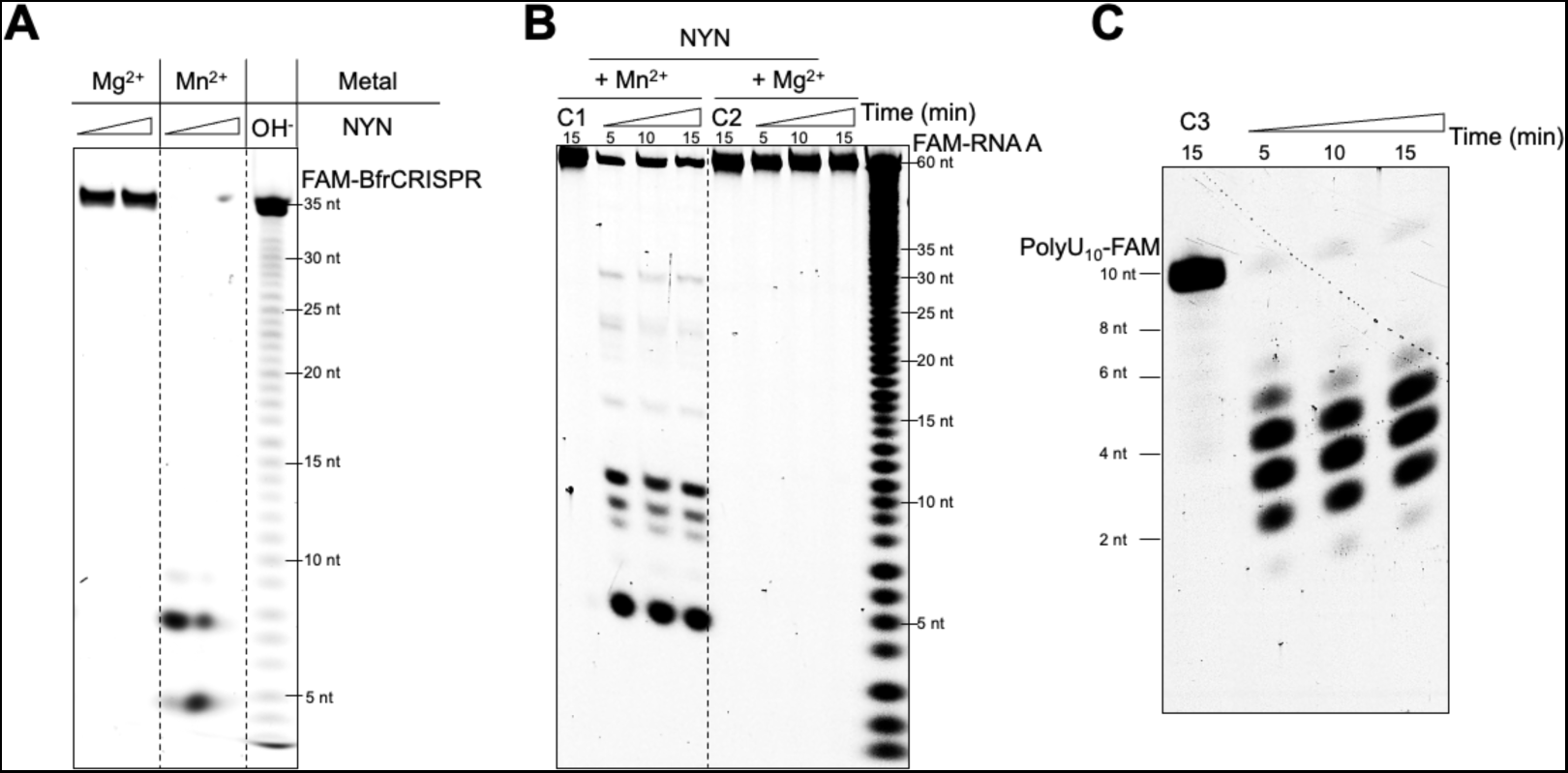
Mn^2+^ dependent endonuclease activity of NYN. **A.** The 5’ FAM labelled RNA BfrCRISPR repeat (35 nt) was incubated with NYN (1 or 5 µM) in the presence of either Mg^2+^ or Mn^2+^ at 37 °C for 1 h. The cleavage activity of NYN was exclusively observed in the presence of Mn^2+^. OH^-^: alkaline lysis marker lane. **B.** The 5’ FAM labelled RNA-A (60 nt) was subjected to incubation with NYN (200 nM) in the presence of Mg^2+^ or Mn^2+^ at 37 °C for 5, 10, or 15 min. Control samples lacking enzymes were included (C1 or C2). Mn^2+^ dependent ribonuclease activity was observed. **C.** The 3’ FAM labelled polyuridine RNA (10 nt) was incubated with NYN (200 nM) in the presence of Mn^2+^ at 37 °C for 5, 10, or 15 min. A control sample lacking NYN was also included. NYN cleaved RNA with a 3’ end label, producing cleavage products without interconversion.

### Predicted structure and active site of *B. fragilis* NYN

We generated a structural model of NYN using Alphafold2 [31] implemented on the Colabfold server [32]. The model, predicted with high confidence, revealed the classic Rossmann fold, characterised by a central 𝛽-sheet comprising six parallel 𝛽-strands, flanked by sets of 𝛼-helices on both sides (Figure 3A, Supplementary Figure 3). Analysis conducted using the DALI server [33] confirmed structural similarities with the uncharacterised NYN-domain protein VPA0982 from *Vibrio parahaemolyticus* (PDB 2QIP; Dali Z-score 17.4; RMSD 2.9 Å over 159 aa) and the N-terminal NYN domain of human MARF1 (PDB 6FDL; Dali Z-score 14.6; RMSD 2.4 Å over 140 aa) [30] (Figure 3B).

**Figure 3.**
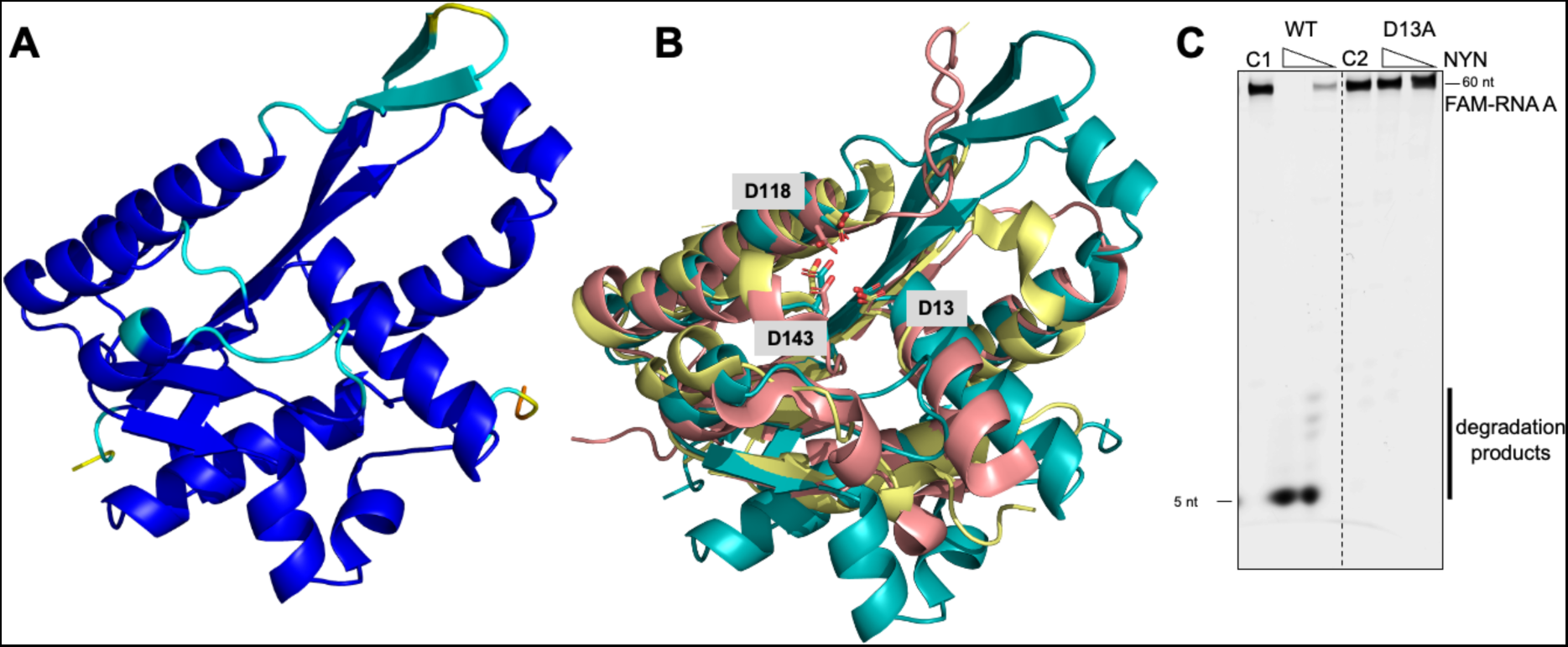
Predicted structure model and mutagenic analysis of the NYN. **A.** A predicted structure model (AlpahFold2) of the NYN. The structural model is shown as a cartoon and prediction confidence (blue to red: high to low confidence) is indicated (predicted local distance difference test (pLDDT)). **B.** Superimposition of the conserved active residues of the NYN (teal) on the VPa0982 (PDB: 2qip, pink) and the human MARF1-NYN domain (PDB: 6fdl, yellow). Predicted active residues D13, D118 and D143 of NYN are highlighted, corresponding to the conserved residues D358, D426, and D452 in MARF1-NYN and the conserved residues D14, D92 and D115 in Vpa0982. **C.** Ribonuclease activity assay of WT and D13A variant NYN. The variant D13A failed to cleave the RNA substrate.

The active sites of NYN domains typically feature a common set of conserved acidic residues [25] which are essential for ribonuclease activity [28]. In the Alphafold2 model of *B. fragilis* NYN, three conserved aspartates (D13, D118 and D143) are clustered in the probable active site of the nuclease (Figure 3B). Structural alignment with VPA0982 (PDB 2QIP) and MARF1-NYN (PDB 6FDL) confirms these 3 residues are absolutely conserved (Figure 3B). To validate these predictions, we mutated *B. fragilis* NYN residue D13 to alanine and assessed the RNA cleavage activity of the variant, which confirmed that residue D13 was indispensable for RNA degradation activity (Figure 3C).

### Potential role of NYN in crRNA maturation

*B. fragilis* NYN functions effectively as a non-specific endoribonuclease *in vitro* and is not essential for Cmr-mediated interference in *E. coli*, but what is its function? One possibility is that NYN plays a role in the maturation of crRNA. Pre-crRNA transcripts are cleaved by Cas6 to generate unit length crRNA species with a long 3’ handle derived from the CRISPR repeat [34, 35]. While these species are incorporated into type I systems where Cas6 is an integral subunit, type III systems process crRNA further, removing a variable amount of the 3’ end, depending on the size of the complex. The Cas6:crRNA intermediate is believed to be transferred to the Csm/Cmr interference complex through a transient interaction [36, 37]. Once bound to the complex, the complex backbone (Csm3/Cmr4) serves as a ruler to determine the length of mature crRNA. Unidentified host nucleases trim exposed 3’ end, leading to crRNA maturation [35, 38–40]. Recent studies have identified host nucleases, including RNase J2, RNase R and PNPase, which promote crRNA maturation in *Staphylococcus epidermidis* type III CRISPR systems [41, 42].

Our previous findings revealed that Cas6 processed the *B. fragilis* CRISPR array within CRISPR repeat as expected, generating a full length, 72 nt crRNA, whereas the mature crRNAs extracted from purified BfrCmr complexes ranged from 37-49 nt in length [23]. This suggests the involvement of unknown ribonucleases in crRNA maturation. We thus hypothesised that NYN might play a role in crRNA maturation in *B. fragilis*. To test this hypothesis, we incubated NYN and Cas6 with an *in vitro* transcribed, radiolabelled mini-CRISPR array (one spacer flanked by two repeats) in the presence or absence of Mn^2+^. Either Cas6 or NYN itself was incubated with this CRISPR array as a control. The reactions were stopped following incubation periods of 5, 10, 30 and 60 min by heating at 95 °C and then analysed on denaturing polyacrylamide gel electrophoresis (PAGE) (Figure 4A). While Cas6 cleaved the CRISPR array as expected, generating a final crRNA of 72 nt, the presence of NYN and Mn^2+^ led to a smear-like degradation pattern of the CRISPR array, consistent with non-specific degradation. The final size of accumulated NYN RNA degradation products was smaller than the mature crRNA (37-49 nt) extracted from Cmr complex, consistent with the absence of Cmr in the reaction (Figure 4 B and C). Additionally, incubation of the purified Cmr complex with NYN had no effect on the size of extracted crRNA, indicating crRNA is protected from nuclease degradation when present in the Cmr complex (Figure 4B).

**Figure 4.**
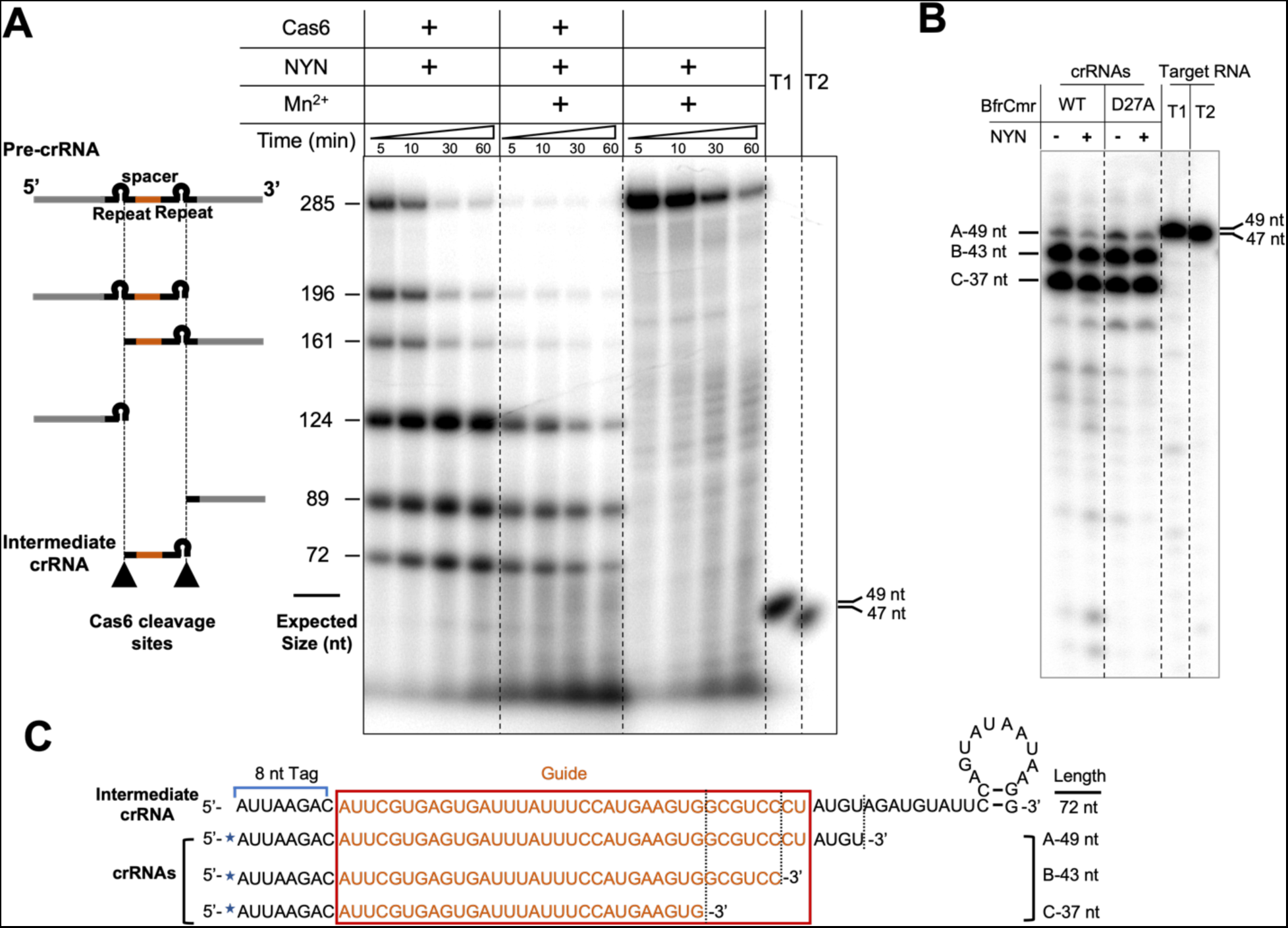
Potential role of NYN in crRNA maturation. **A.** Ribonuclease activity assay of NYN towards precursor crRNA (pre-crRNA). An internally radio-labelled transcript RNA containing two CRISPR repeats (black) and one spacer sequence targeting Phage P1 (orange) was incubated with Cas6 (1 μM) and/or NYN (70 nM) for the indicated times and analysed by denaturing gel electrophoresis. The expected sizes and compositions of cleavage products of Cas6 are indicated based on its specific cleavage sites within each repeat (indicated by black triangles). Radio-labelled target RNA T1 (Target RNA_Lpa, 49 nt) and T2 (Non-target RNA_pUC, 47 nt) were used as size markers. **B.** Comparison of the size of crRNAs extracted from purified wild type or Cmr4 D27A variant *B. fragilis* Cmr complex, in the presence or absence of NYN. Radio-labelled target RNAs T1 and T2 were used to indicate the size. **C.** Schematic comparison of sequence and size of an intermediate crRNA and extracted Cmr-bound crRNA species. An intermediate crRNA, derived from Cas6 cleavage, contains the 5’ end repeat-derived tag, spacer sequence (centre) and 3’ end repeat-derived sequence with a stem-loop. Extracted crRNA species are all shorter than an intermediate crRNA, indicating trimming from the 3’ end. Extracted crRNAs were labelled at the 5’ end with ^32^ P (blue star).

We observed that Cas6 initially co-purifies with the *B. fragilis* Cmr complex when they are co-expressed in *E. coli*, suggestive an interaction that is lost in the final stages of purification (Supplementary Figure 4). A possible explanation for this is that Cas6 remains bound to the crRNA after cleaving it and is only separated from the complex when the crRNA is further processed. To test this, we incubated Cas6 with the FAM-labelled *B. fragilis* CRISPR repeat *in vitro*, then analysed the products by electrophoretic mobility shift assay (EMSA). Under these conditions, Cas6 was expected to cleave the CRISPR RNA. As we increased the concentration of Cas6, we observed the progressive accumulation of a retarded RNA species corresponding to a Cas6:CRISPR RNA species (Figure 5A,B), confirming formation of a stable complex. To investigate whether the NYN protein could cleave this Cas6:CRISPR RNA complex, we incubated the FAM-labelled CRISPR RNA with Cas6, NYN or both proteins together, then analysed the reaction products (Figure 5C). As expected, Cas6 efficiently cleaved the CRISPR RNA at the base of the hairpin, while NYN alone fully degraded the CRISPR RNA to small products. When both proteins were present, NYN could still cleave the CRISPR RNA near the 5’ end, but a significant fraction of the RNA was protected from cleavage. This confirms that RNA binding proteins can protect bound RNA from NYN cleavage – as observed for many other non-specific nucleases. Overall, these biochemical data are consistent with a role for NYN in crRNA maturation.

**Figure 5.**
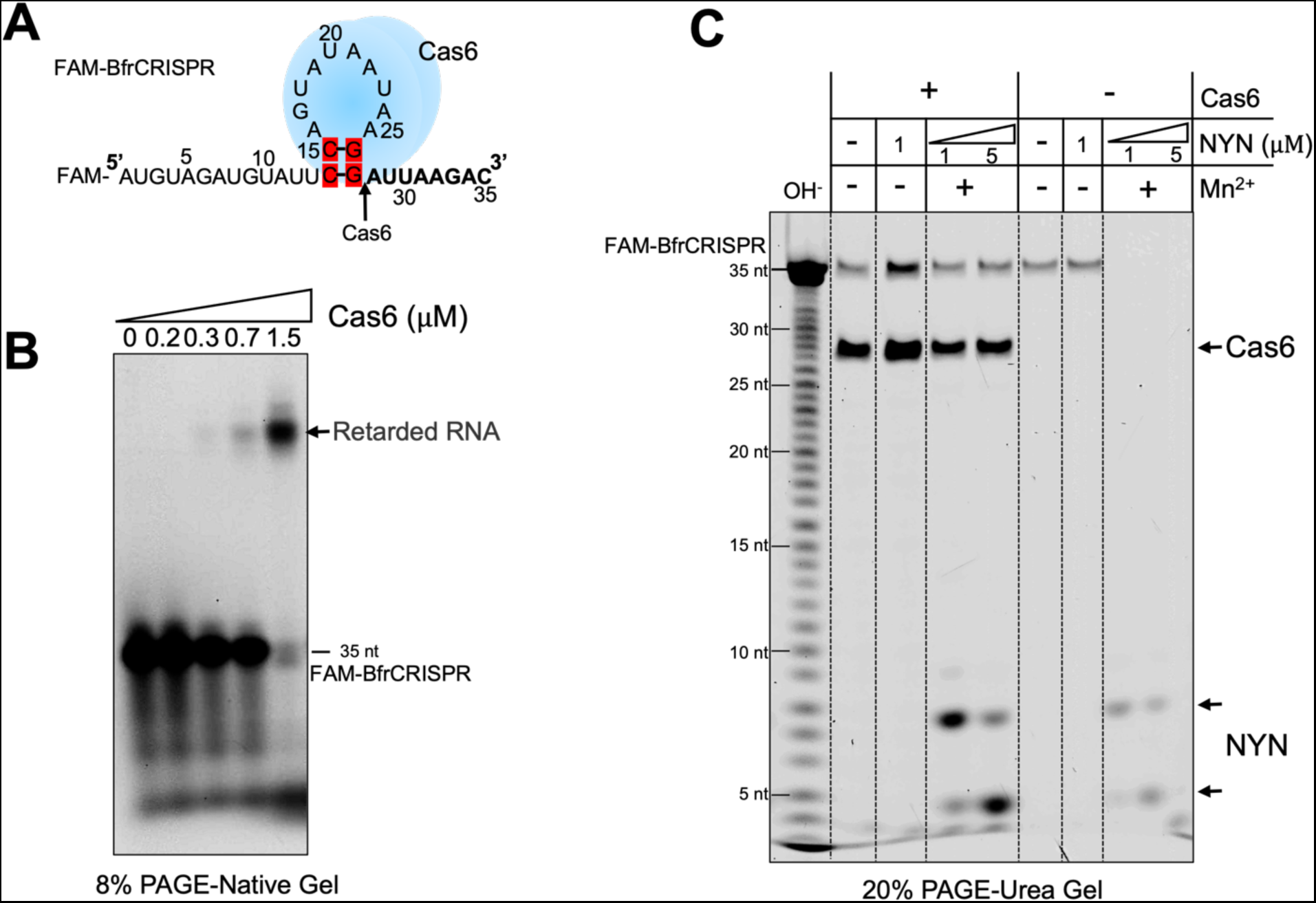
Ribonuclease activity of NYN in the presence of Cas6. **A.** Predicted secondary structure of BfrCRISPR repeat RNA. Cas6 (blue) is predicted to bind to the stem loop of CRISPR repeat RNA, with its cleavage site indicated by a black arrow. **B.** Cas6 binds CRISPR repeat RNA. 5’-FAM labelled CRISPR repeat RNA (100 nM) was incubated with varying concentrations of Cas6 (0, 0.2, 0.3, 0.7, 1.5 μM) at 37 °C for 1 h. The appearance of a retarded RNA species was observed. **C.** Bound RNA is protected from NYN cleavage. 5’-FAM labelled CRISPR repeat RNA (100 nM) was incubated with NYN (1 or 5 μM) in the presence or absence of Cas6 (3 μM) at 37°C for 1 h. NYN cleavage activity is hindered when in the presence of Cas6, suggesting the bound RNA is protected from NYN-mediated cleavage. OH^-^: alkaline lysis marker lane. The cleavage products of Cas6 and NYN are indicated at the side of the gel.

## Discussion

Type III CRISPR systems are widespread in prokaryotes and exhibit considerable diversity in their components and mechanisms for anti-viral immunity. Here, we addressed the function of the CRISPR-associated NYN ribonuclease, which is found associated with a subset of type III-B CRISPR systems [22], including the SAM-AMP signalling system from *B. fragilis* [23]. Unlike the cyclic oligoadenylate-activated ribonucleases Csm6 and Csx1, NYN does not appear to function directly in anti-viral defence [23].

Type III CRISPR systems require a multi-stage crRNA processing pathway initiated by Cas6-mediated cleavage of the CRISPR repeat, followed by trimming of the 3’ end. Type III CRISPR effectors commonly vary in size by one Cas7/Cas11 backbone “unit”, coinciding with a range of bound crRNAs varying in size by 6 nt [35, 39, 43, 44]. This is consistent with footprinting, or protection, of bound crRNA, leaving an exposed 3’ RNA end that is trimmed by ribonucleases. Indeed, elegant studies in *Staphylococcus epidermidis* have demonstrated that a range of ribonucleases including RNase J2, PNPase and RNase R are responsible for this role [41, 42]. These enzymes are widely conserved with homologues present in *E. coli* and *B. fragilis* and have many housekeeping duties in the cell relating to RNA metabolism. To ensure efficient recruitment to crRNA processing in *S. epidermidis*, RNase R has been shown to interact directly with the Cmr5 subunit of the type III CRISPR system [41].

Here, we have demonstrated that NYN is a Mn-dependent, non-sequence specific ribonuclease, which can cleave RNA internally and is capable of cutting near 5’ and 3’ termini. NYN is also sensitive to footprinting by ribonucleoprotein complexes, and efficiently cleaves the *B. fragilis* CRISPR RNA when it is exposed. These properties, coupled with the location of the *nyn* gene next to *cas6*, prompt us to propose a function for CRISPR-associated NYN as a dedicated crRNA 3’ end trimming ribonuclease (Figure 6). There are several limitations to our study. Firstly, we have not directly tested for crRNA processing in the presence of the apo-Cmr complex *in vitro*. These are challenging experiments requiring expression of all six Cmr subunits in the absence of crRNA. Secondly, we have been unable to explore the importance of NYN in *B. fragilis*. We postulate that NYN is not required for type III CRISPR immunity in the heterologous context in *E. coli* due to the availability of alternative RNA processing enzymes. Finally, we cannot rule out a role for NYN in the generation of spacers for CRISPR adaptation. These originate as RNA species and are converted to DNA by the RT-Cas1 enzyme prior to incorporation in the CRISPR locus [45]. Clearly, NYN could conceivably play a role in this process, for example by trimming RNA protospacers. This possibility warrants further investigation but is beyond the scope of this study.

**Figure 6.**
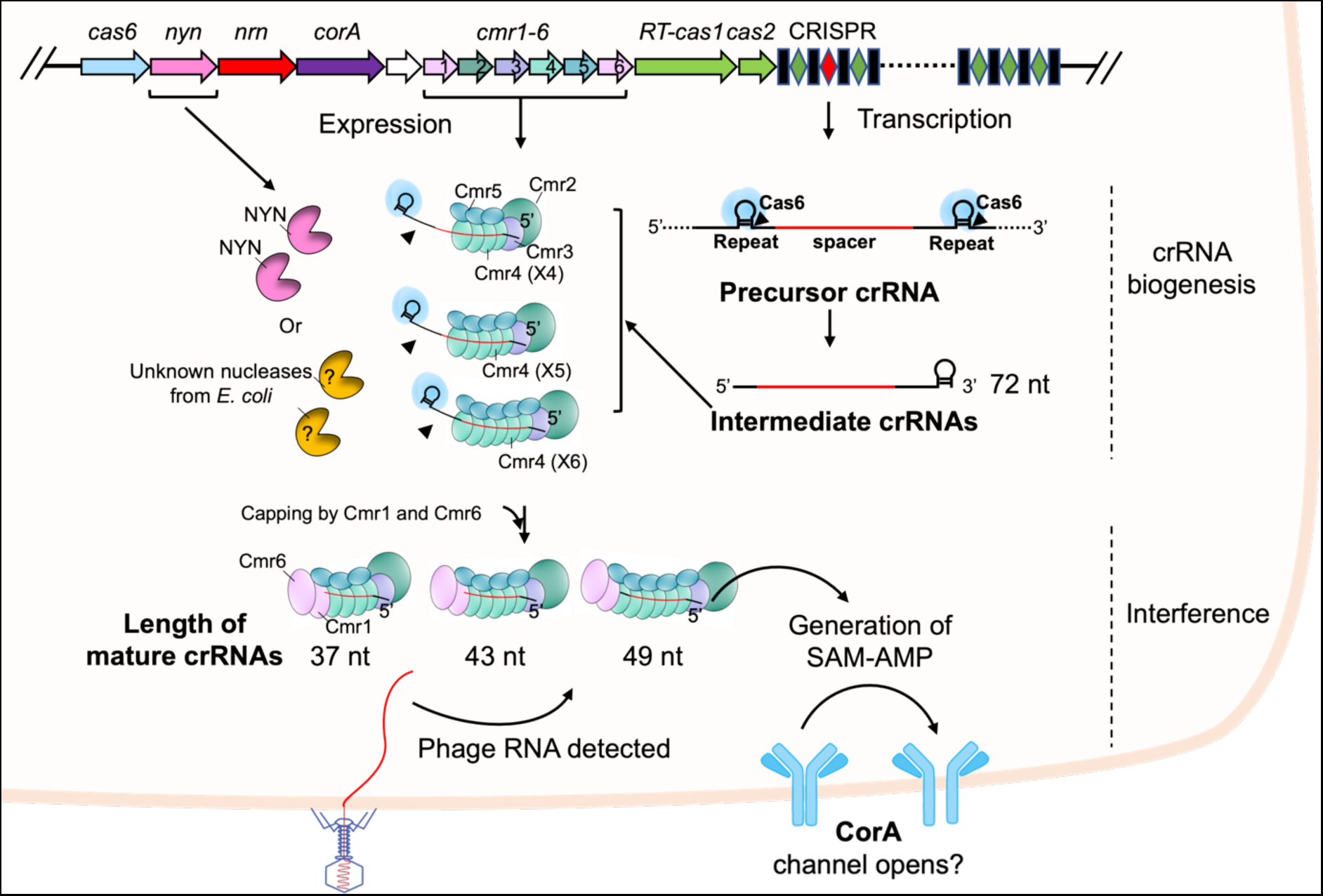
Model of *B. fragilis* crRNA maturation. In the crRNA biogenesis stage, the CRISPR array undergoes transcription into precursor crRNA, which is subsequently processed by Cas6 into intermediate crRNAs (72 nt) at the base of a predicted hairpin structure. The BfrCmr subunits are assembled around intermediate RNAs in various stoichiometries. The exposed regions of Cmr-bound intermediate crRNAs are further trimmed by endonuclease NYN in *B. fragilis* or unknown nucleases in *E. coli*. Following this processing, subunits Cmr1 and Cmr6 are capped at the 3’ end of crRNA. The length of matured crRNAs (37, 43 and 49 nt) is determined by the number of backbone subunits Cmr4 (Cas7) and Cmr5 (Cas11).

## Materials and Methods

### Cloning

The gene encoding *B. fragilis* NYN was codon optimised and synthesised for expression in *Escherichia coli*. A synthetic g-block was purchased from Integrated DNA Technologies (IDT) and cloned into plasmid pEhisV5Tev [46] between the *Nco*I and *Bam*HI sites. Plasmid construction was performed in *E. coli* DH5α, and sequence integrity was confirmed by sequencing (Eurofins Genomics). *E. coli* C43 (DE3) cells were transformed with the verified plasmid for protein expression. Plasmid pRATDuet-NYN used in plasmid challenge assay was constructed by inserting the *B. fragilis nyn* gene between *Nco*I and *EcoR*I restriction sites of pRATDuet [19] under control of pBAD promoter.

### Protein expression and Purification

*E. coli* C43 (DE3) cells containing the verified plasmid were grown for 18 h at 16 °C with shaking after induction with 0.2 mM isopropyl β-D-1-thiogalactopyranoside (IPTG) at an OD_600_ of 0.6∼0.8. Cell pellets were harvested by centrifugation at 5000 rpm (Beckman Coulter Avanti JXN-26; JLA8.1 rotor) at 4 °C for 15 min. Cell pellets were resuspended into 4 volumes equivalent of lysis buffer (50 mM Tris-HCl pH 8.0, 0.5 M NaCl, 10 mM imidazole, and 10% glycerol) supplemented with lysozyme (1 mg/ml) and EDTA-free protease inhibitor tablets (Roche). Pellets were lysed by sonicating for 6 min with 1 min rest intervals on ice, before pelleting cell debris by ultracentrifugation at 40,000 rpm (Beckman Coulter Optima L-90K, 70 Ti rotor) at 4 °C for 30 min.

Supernatants were subsequently loaded onto a 5 mL HisTrap FF column (GE Healthcare) equilibrated with lysis buffer. After washing away unbound proteins with 20 column volumes of lysis buffer, the his-tagged protein was eluted with linear gradient elution of elution buffer (50 mM Tris-HCl pH 8.0, 0.5 M NaCl, 0.5 M imidazole, and 10% glycerol). Fractions containing recombinant protein were collected and concentrated for his-tag removal by overnight dialysis with TEV protease (1 mg per 10 mg protein) in size exclusion chromatography (SEC) buffer (20 mM Tris-HCl pH 8.0, 0.25 M NaCl, 1 mM DTT, and 10% glycerol). The TEV-cleaved protein was recovered by second immobilised metal affinity chromatography (IMAC) and further purified by SEC (S200 16/60 column, GE Healthcare) in SEC buffer under isocratic flow. The purified protein was flash frozen and stored at −70 °C. SDS-PAGE was performed at each purification step to evaluate purity and integrity of the protein.

### Ribonuclease assay

The ribonuclease activity of NYN was conducted by incubating the indicated concentration of NYN protein with 400 nM RNA (Table 1) with a 5’ or 3’ fluorescein (FAM) label, in the buffer of 20 mM Tris-HCl, pH 7.5, 50 mM NaCl and 5 mM MnCl_2_ (or MgCl_2_) at 37 °C for 1 h. For assays analysis crRNA transcript cleavage, 70 nM NYN and/or 1 μM Cas6 were incubated with an internally radio-labelled transcript RNA (100 nM) (Table 1) in the same buffer condition at 37 °C for 5, 10, 30 and 60 min. 5’-FAM labelled CRISPR repeat RNA (100 nM) was incubated with 1 or 5 μM NYN in the presence or absence of 3 μM Cas6 at 37°C for 1 h or time courses assay by incubating 500 nM NYN with or without 1 μM Cas6 for 5, 15, 30 and 60 min. All reactions were stopped by adding10 mM EDTA, before heating at 95 °C for 10 min, followed by incubation on ice.

**Table 1.**
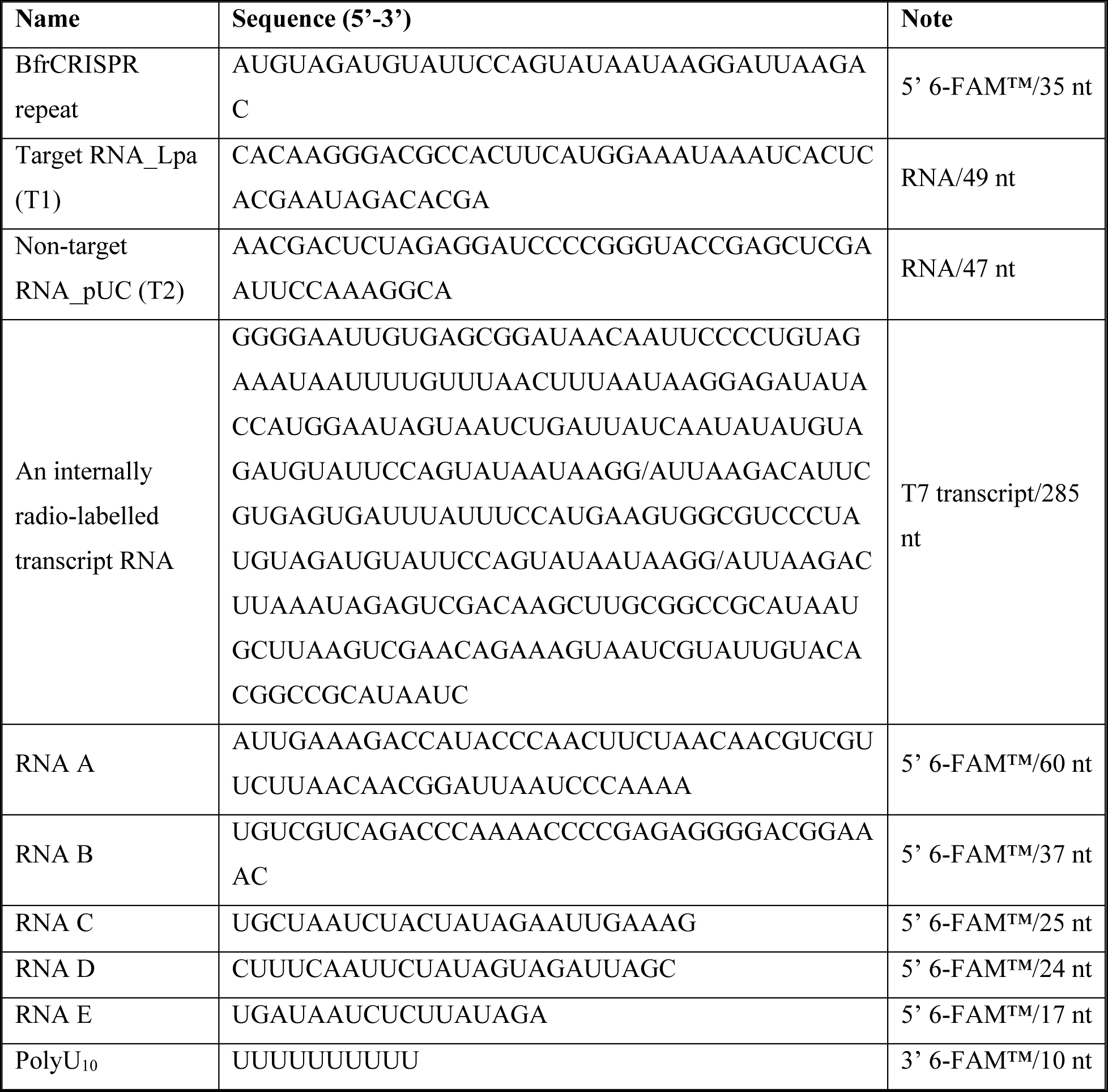
RNA sequences used in this study.

All samples were analysed on 20% acrylamide, 7 M urea and 1X TBE denaturing gels, which were run at 30 W and 45 °C for 2 h. The gel was finally imaged by Typhoon FLA 7000 imager (GE Healthcare) at a wavelength of 650 nm for phosphoimaging and 473 nm for FAM fluorescence (pmt 700∼900). An alkaline hydrolysis ladder used for cleavage site mapping was generated by incubating RNA substrates in a buffer of 5 mM NaHCO_3_, pH 9.5 at 95 °C for 5 min.

### Electrophoretic mobility shift assay

100 nM 5’-FAM labelled CRISPR repeat RNA was subjected to incubation with varying concentrations of purified Cas6 (0, 0.2, 0.3, 0.7, 1.5 μM) in buffer (20 mM Tris-HCl, pH 7.5, 50 mM NaCl and 1 mM EDTA) at 37 ℃ for 1 h. Reactions were mixed with ficoll loading buffer and then analysed on the native acrylamide gel (8% (w/v) 19:1 acrylamide: bis-acrylamide). Electrophoresis was performed in 1X TBE running buffer at 200V for 2h at room temperature. The gel was imaged using a Typhoon FLA 7000 imager (GE Healthcare) at a wavelength of 473 nm for FAM fluorescence (pmt 700–900).

### Production of radio-labelled transcripts

The plasmid pBfrCRISPR_Lpa was described previously [23]. An internally radio-labelled transcript RNA containing two BfrCRISPR repeats, and one guide sequence (Table 1) was generated by following the MEGAscript®Kit (Invitrogen) protocol. The template used in transcription was obtained by PCR of plasmid pBfrCRISPR_Lpa using primer Duet-up and Duet-Down. 120 ng PCR product was mixed with 7.5 mM ribonucleotide solution (ATP, GTP, UTP, CTP) and 133 nM α-^32^P-ATP (Perkin Elmer) as a tracer, before incubation at 37 °C for 4 h in the 1X reaction buffer with T7 enzyme mix. Transcript was subsequently purified by phenol: chloroform extraction and isopropanol precipitation. The purity of transcript was check on urea polyacrylamide denaturing gel.

### Generation of 5’ end radio-labelled RNAs

RNAs (10-50 μM) with a 5’ terminal hydroxyl group were incubated with T4 polynucleotide kinase (Thermo Scientific) and 𝛾-^32^P-ATP (10 mCi/ml, Perkin Elmer) at 37 °C for 40 min, before quenching with 25 mM EDTA. The radio-labelled RNAs Target_Lpa and Target_pUC (Table 1) or extracted crRNAs were subsequently loaded and purified on a 20% denaturing polyacrylamide gel (7 M urea and 1X TBE) at 30 W and 45 °C for 2 h. The band containing radio-labelled RNAs was excised from the gel, crushed and then soaked into 500 μl RNase-free H_2_O overnight at 4 °C. Gel particles were removed by centrifugation at 5,000 rpm at 4 °C for 30 s and the radio-labelled RNAs from supernatants were subsequently recovered by phenol: chloroform extraction and isopropanol precipitation.

### crRNA recovery from the *B. fragilis* Cmr complex

crRNAs from the purified Cmr complex were all recovered by phenol: chloroform extraction and isopropanol precipitation. Samples were mixed thoroughly with ammonium acetate to a final concentration of 0.5 M, before extraction with an equal volume of phenol: chloroform and then with an equal volume of chloroform. RNAs in the aqueous phase were precipitated by mixing with one volume of isopropanol. The mixture was chilled at −20 °C for at least 15 min. RNAs were pelleted by centrifugation at 4 °C for 15 min at maximum speed and subsequently resuspended in RNase-free H_2_O, after removing the supernatant solution.

### SAM-AMP cleavage activity

The methods to generate SAM-AMP *in vivo* and *in vitro* were described previously [23]. Briefly, a single colony of *E. coli* BL21star containing the plasmids pBfrCmr1-6, pBfrCRISPR_Tet (or pBfrCRISPR_pUC) and pRATDuet-NYN (or pRATDuet) was inoculated into 10 mL of L-broth (LB) with antibiotic (50 µg/ml ampicillin, 50 µg/ml spectinomycin and 12.5 µg/ml tetracycline). Cells were grown overnight at 37 °C with shaking at 180 rpm, before recultivating with 20-fold dilution into 20 ml fresh LB with same antibiotics and continually incubating at 37 °C. The cell culture was fully induced by adding 0.2 % (w/v) D-lactose and 0.2 % (w/v) L-arabinose at OD_600_ of 0.4 - 0.6. After overnight induction at 25 °C, the cell culture was mixed with 4 volumes equivalent of cold PBS and pelleted by centrifugation at 4,000 rpm at 4 °C for 10 min. Pellets were resuspended into 2 ml cold extraction solvent [acetonitrile/methanol/water (2/2/1, v/v/v)], vortexed for 30 s and stored at −20 °C overnight. Supernatant was obtained by centrifugation at 4 °C for 15 min at maximum speed, followed by HPLC analysis.

Purified NYN (5 µM) was incubated with SAM-AMP (100 µM) in the buffer of 20 mM Tris-HCl, pH 7.5, 50 mM NaCl and 5 mM MnCl_2_ at 37 °C for 1 h. The reaction was quenched by adding EDTA to a final concentration of 50 mM. Samples were purified by microfiltration (Pall Nanosep® Spin Filter, MWCO 3 kDa) and subjected to HPLC analysis.

### HPLC analysis

HPLC analysis was described previously [23]. In brief, sample analysis was conducted on an UltiMate 3000 UHPLC system (ThermoFisher scientific) with absorbance monitoring at 260 nm. Samples were injected into a C18 column (Kinetex EVO 2.1 X 50 mm, the particle size of 2.6 µm) at 40 °C and analysed with gradient elution in solvent A (10 mM ammonium bicarbonate) and solvent B (Acetonitrile with 0.1 % TFA) at a flow rate of 0.3 ml/min.

## Acknowledgements

We thank Dr Sabine Grüschow and Dr Ville Hoikkala for help and advice.

## Data Availability

All supporting data are included within the main article and its supplementary files.

## Supplementary Figures

**Supplementary Figure 1.**
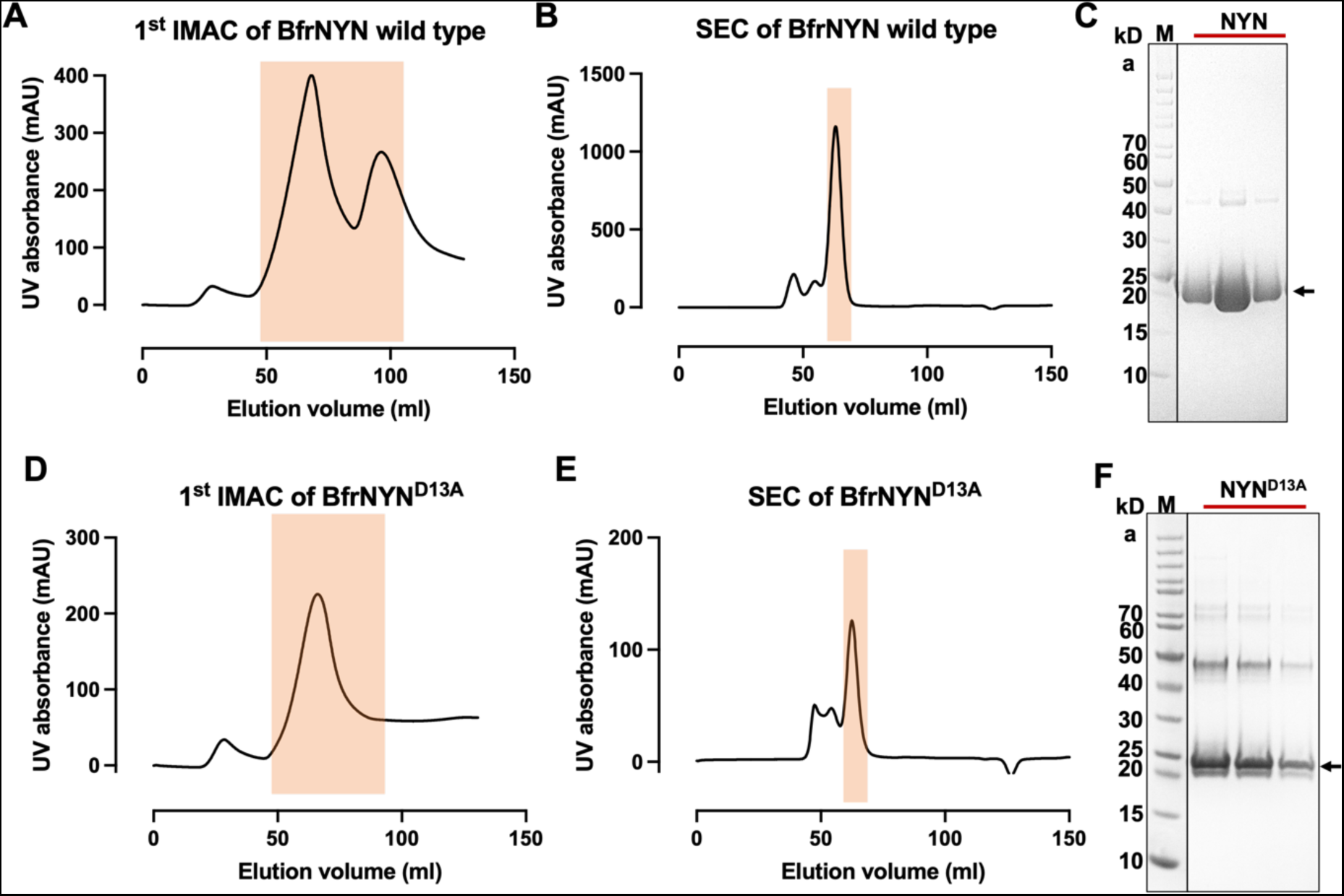
The purification of NYN wild type and variant D13A. **A, and D.** The first immobilised metal affinity chromatography (1st IMAC) step of NYN wild type and variant D13A. The shaded fractions containing target protein were pooled for his tag removal. **B, and E.** Superdex200 SEC profiles. The TEV-cleaved protein was recovered from the nickel column and then subjected to SEC. Shaded fractions were collected and concentrated for further enzymatic analysis. **C, and F.** SDS-PAGE analysis of purity of NYN wild type and variant D13A. The monomer mass is approximately 25 kDa, consistent with the theoretical mass of NYN. M is the marker to indicate size on the gel. Some dimeric species (48 kDa) were observed in the SDS-PAGE.

**Supplementary Figure 2.**
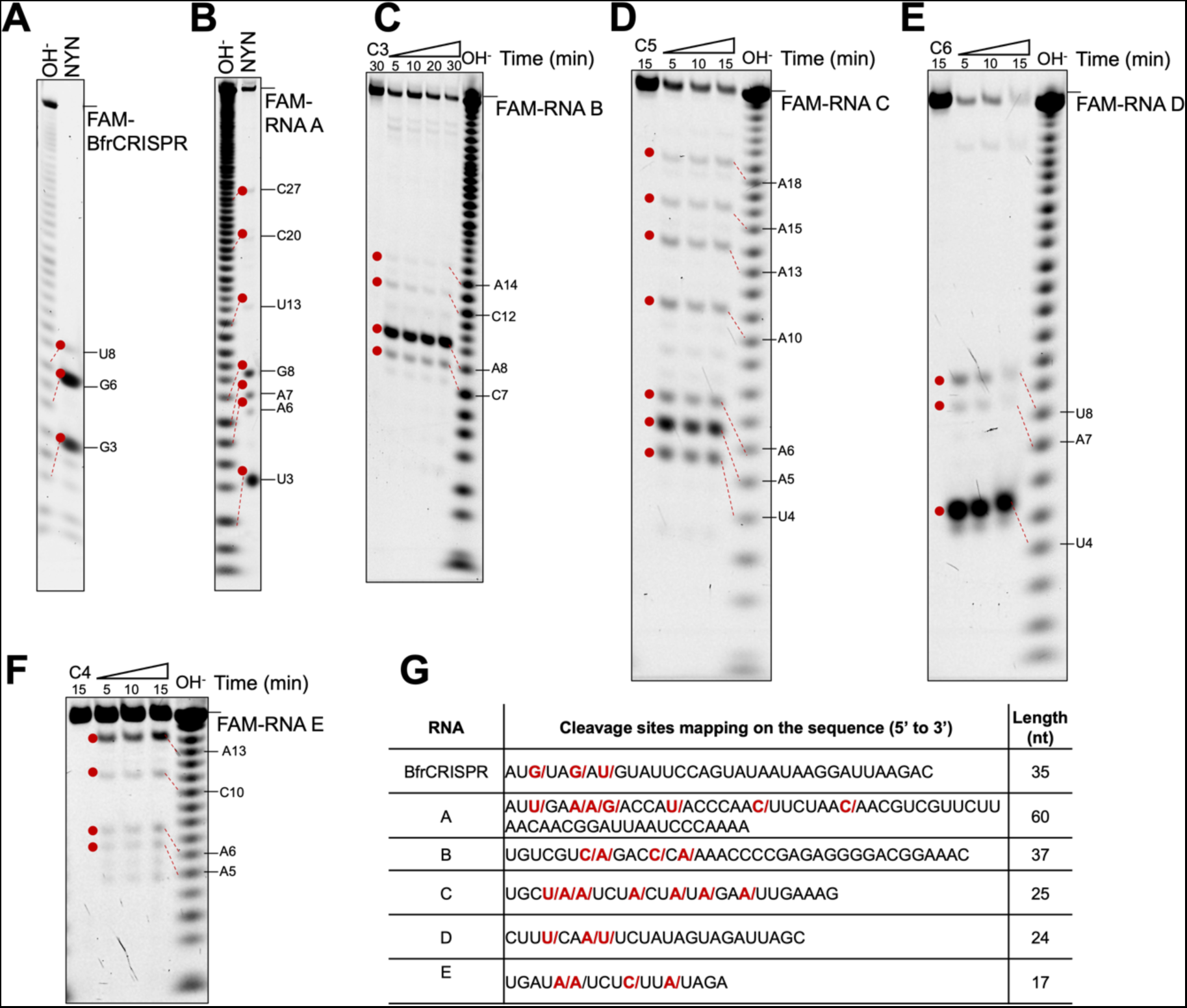
Mapping the RNA cleavage sites of NYN. **A-F.** Mapping of NYN cleavage sites towards various RNA substrates. 5’ FAM labelled RNA-BfrCRISPR, A, B, C, D and E were incubated with 200 nM NYN in the presence of Mn^2+^ at 37°C for 30 min or indicated time points. Cleavage sites were mapped against an alkaline hydrolysis ladder of corresponding RNA, indicated by red dashed lines. Sequence assignments of bands are shown along the side of the gel. The cleaved species indicated by red dots migrate one and half nucleotides slower than bands in the hydrolysis ladder [47], given that the alkaline hydrolysis RNA harbouring a phosphate group at 3’ end [48] and Mn^2+^ dependent NYN cleavage products leaving a hydroxyl group at 3’end [49]. **G.** Table of mapped cleavage sites. The cleavage sites were highlighted with red slash and nucleotides indicated alongside the gel were in bold and red.

**Supplementary Figure 3.**
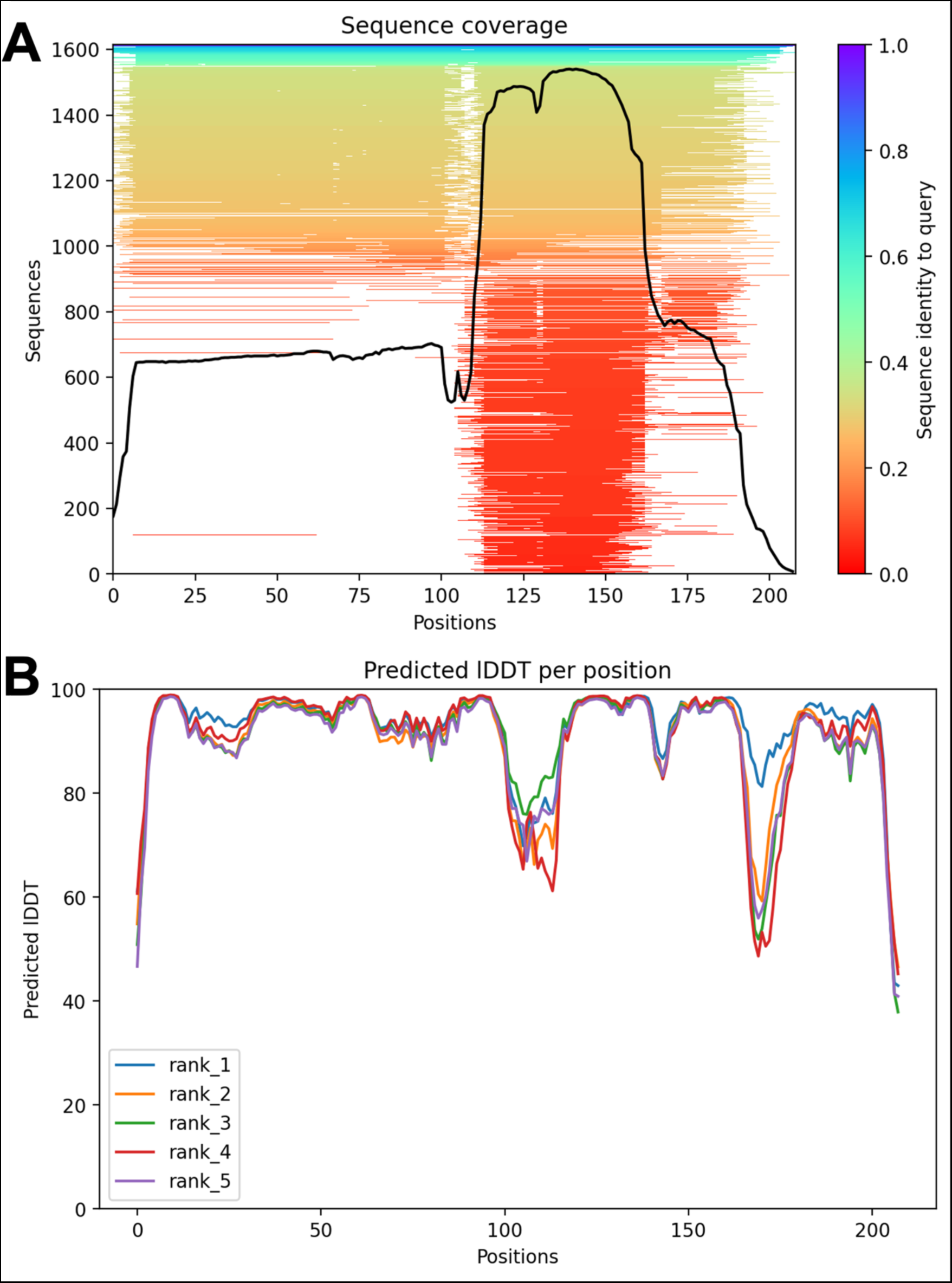
Sequence coverage and local distance difference test (IDDT) scores for the Alphafold2 model of NYN.

**Supplementary Figure 4.**
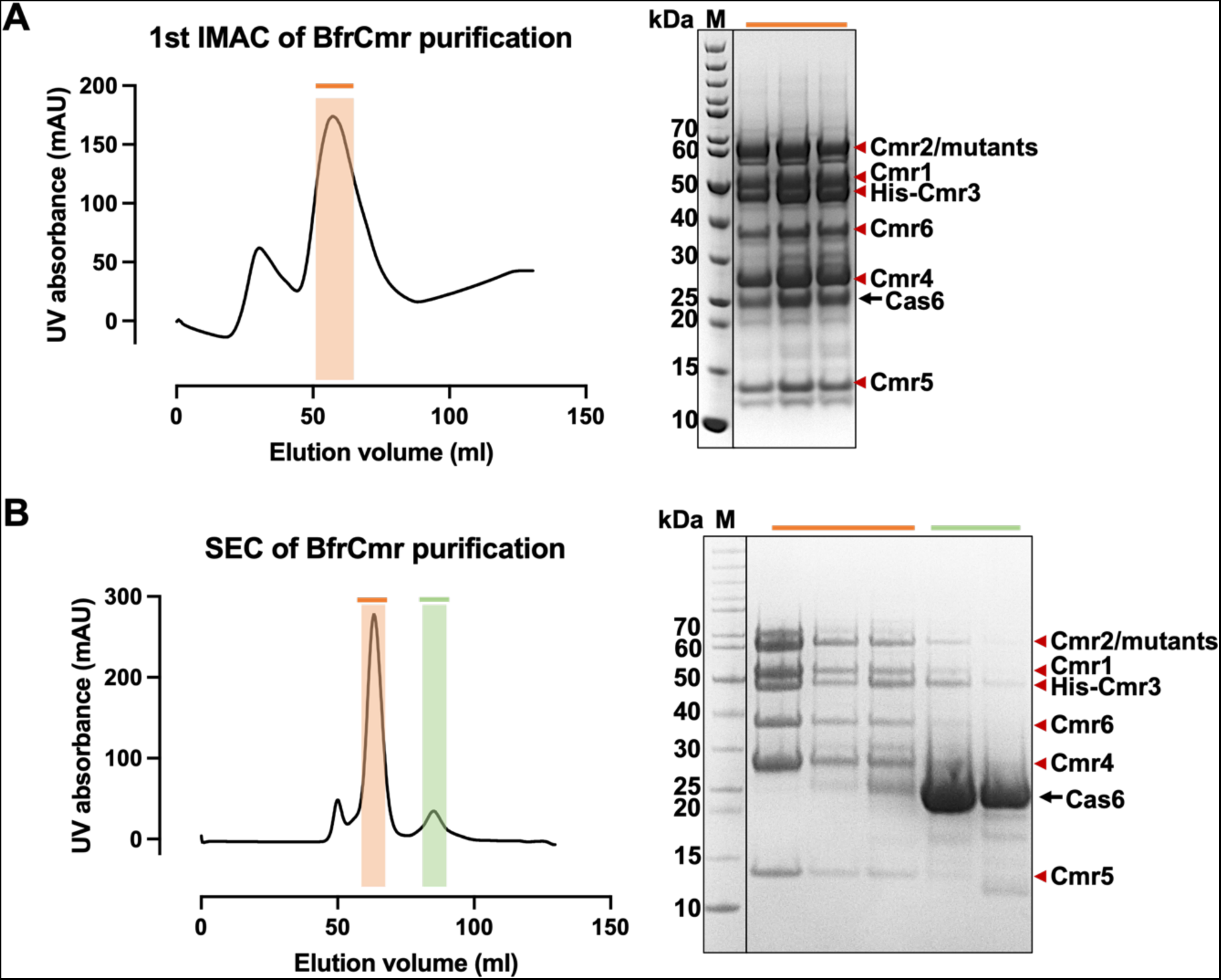
Cas6 co-purifies with the BfrCmr complex. **A.** The first immobilised metal affinity chromatography (1st IMAC) of BfrCmr complex. The shaded fractions were analysed by SDS-PAGE. Cas6 was co-purified with Cmr complex. **B.** Superdex200 SEC profiles. Fractions pooled from 1^st^ IMAC was subjected to SEC. Shaded fractions were analysed by SDS-PAGE. Cas6 was separated from BfrCmr complex during SEC. The monomer mass of Cas6 is approximately 26 kDa. M: protein size standards.

## References

1 Gasiunas, G., Sinkunas, T. and Siksnys, V. (2014) Molecular mechanisms of CRISPR-mediated microbial immunity. Cell Mol Life Sci. 71, 449–465

2 Hille, F., Richter, H., Wong, S. P., Bratovic, M., Ressel, S. and Charpentier, E. (2018) The Biology of CRISPR-Cas: Backward and Forward. Cell. 172, 1239–1259

3 Sasnauskas, G. and Siksnys, V. (2020) CRISPR adaptation from a structural perspective. Curr Opin Struct Biol. 65, 17–25

4 Levy, A., Goren, M. G., Yosef, I., Auster, O., Manor, M., Amitai, G., Edgar, R., Qimron, U. and Sorek, R. (2015) CRISPR adaptation biases explain preference for acquisition of foreign DNA. Nature. 520, 505–510

5 Charpentier, E., van der Oost, J., White, M.F. (2015) Biogenesis pathways of RNA guides in archaeal and bacterial CRISPR-Cas adaptive immunity. FEMS Microbiol Rev, 39:428–441

6 Nussenzweig, P. M. and Marraffini, L. A. (2020) Molecular Mechanisms of CRISPR-Cas Immunity in Bacteria. In Annual Review of Genetics, 54, 2020 (Bonini, N. M., ed.). pp. 93–120

7 Makarova, K. S., Wolf, Y. I., Iranzo, J., Shmakov, S. A., Alkhnbashi, O. S., Brouns, S. J. J., Charpentier, E., Cheng, D., Haft, D. H., Horvath, P., Moineau, S., Mojica, F. J. M., Scott, D., Shah, S. A., Siksnys, V., Terns, M. P., Venclovas, C., White, M. F., Yakunin, A. F., Yan, W., Zhang, F., Garrett, R. A., Backofen, R., van der Oost, J., Barrangou, R. and Koonin, E. V. (2020) Evolutionary classification of CRISPR-Cas systems: a burst of class 2 and derived variants. Nat Rev Microbiol. 18, 67–83

8 Koonin, E. V., Makarova, K. S. and Zhang, F. (2017) Diversity, classification and evolution of CRISPR-Cas systems. Curr Opin Microbiol. 37, 67–78

9 Makarova, K. S. (2015) An updated evolutionary classification of CRISPR–Cas systems. Nat. Rev. Microbiol. 13: 722–736

10 Athukoralage, J. S. and White, M. F. (2021) Cyclic oligoadenylate signalling and regulation by ring nucleases during type III CRISPR defence. RNA. 27, 855–867

11 Niewoehner, O. (2017) Type III CRISPR–Cas systems produce cyclic oligoadenylate second messengers. Nature. 548:543–548

12 Kazlauskiene, M., Kostiuk, G., Venclovas, Č., Tamulaitis, G. and Siksnys, V. (2017) A cyclic oligonucleotide signaling pathway in type III CRISPR-Cas systems. Science. 357:605–609

13 Meeske, A. J., Nakandakari-Higa, S. and Marraffini, L. A. (2019) Cas13-induced cellular dormancy prevents the rise of CRISPR-resistant bacteriophage. Nature. 570, 241–245

14 Athukoralage, J. S., Graham, S., Gruschow, S., Rouillon, C. and White, M. F. (2019) A Type III CRISPR Ancillary Ribonuclease Degrades Its Cyclic Oligoadenylate Activator. Journal of Molecular Biology. 431, 2894–2899

15 Athukoralage, J. S., Rouillon, C., Graham, S., Gruschow, S. and White, M. F. (2018) Ring nucleases deactivate type III CRISPR ribonucleases by degrading cyclic oligoadenylate. Nature. 562, 277–280

16 Jia, N., Jones, R., Yang, G., Ouerfelli, O. and Patel, D. J. (2019) CRISPR-Cas III-A Csm6 CARF Domain Is a Ring Nuclease Triggering Stepwise cA(4) Cleavage with ApA > p Formation Terminating RNase Activity. Molecular Cell. 75, 944–956

17 Garcia-Doval, C., Schwede, F., Berk, C., Rostol, J. T., Niewoehner, O., Tejero, O., Hall, J., Marraffini, L. A. and Jinek, M. (2020) Activation and self-inactivation mechanisms of the cyclic oligoadenylate-dependent CRISPR ribonuclease Csm6. Nature Communications. 11:1596

18 Smalakyte, D., Kazlauskiene, M., Havelund, J. F., Ruksenaite, A., Rimaite, A., Tamulaitiene, G., Faergeman, N. J., Tamulaitis, G. and Siksnys, V. (2020) Type III-A CRISPR-associated protein Csm6 degrades cyclic hexa-adenylate activator using both CARF and HEPN domains. Nucleic Acids Research. 48, 9204–9217

19 Athukoralage, J. S., McMahon, S. A., Zhang, C., Gruschow, S., Graham, S., Krupovic, M., Whitaker, R. J., Gloster, T. M. and White, M. F. (2020) An anti-CRISPR viral ring nuclease subverts type III CRISPR immunity. Nature. 577, 572–575

20 Athukoralage, J. S., Graham, S., Rouillon, C., Gruschow, S., Czekster, C. M. and White, M. F. (2020) The dynamic interplay of host and viral enzymes in type III CRISPR-mediated cyclic nucleotide signalling. Elife. 9, e55852

21 Schmier, B. J., Nelersa, C. M. and Malhotra, A. (2017) Structural Basis for the Bidirectional Activity of Bacillus nanoRNase NrnA. Sci Rep. 7, 11085

22 Shmakov, S. A., Makarova, K. S., Wolf, Y. I., Severinov, K. V. and Koonin, E. V. (2018) Systematic prediction of genes functionally linked to CRISPR-Cas systems by gene neighborhood analysis. Proc Natl Acad Sci U S A. 115, E5307–e5316

23 Chi, H., Hoikkala, V., Gruschow, S., Graham, S., Shirran, S. and White, M. F. (2023) Antiviral type III CRISPR signalling via conjugation of ATP and SAM. Nature. 622, 826–833

24 Faure, G., Makarova, K. S. and Koonin, E. V. (2019) CRISPR-Cas: Complex Functional Networks and Multiple Roles beyond Adaptive Immunity. J Mol Biol. 431, 3–20

25 Anantharaman, V. and Aravind, L. (2006) The NYN domains: novel predicted RNAses with a PIN domain-like fold. RNA Biol. 3, 18–27

26 Matelska, D., Steczkiewicz, K. and Ginalski, K. (2017) Comprehensive classification of the PIN domain-like superfamily. Nucleic Acids Res. 45, 6995–7020

27 Leroy, M., Piton, J., Gilet, L., Pellegrini, O., Proux, C., Coppée, J. Y., Figaro, S. and Condon, C. (2017) Rae1/YacP, a new endoribonuclease involved in ribosome-dependent mRNA decay in Bacillus subtilis. Embo j. 36, 1167–1181

28 Yao, Q., Cao, G., Li, M., Wu, B., Zhang, X., Zhang, T., Guo, J., Yin, H., Shi, L., Chen, J., Yu, X., Zheng, L., Ma, J. and Su, Y. Q. (2018) Ribonuclease activity of MARF1 controls oocyte RNA homeostasis and genome integrity in mice. Proc Natl Acad Sci U S A. 115, 11250–11255

29 Faure, G., Shmakov, S. A., Yan, W. X., Cheng, D. R., Scott, D. A., Peters, J. E., Makarova, K. S. and Koonin, E. V. (2019) CRISPR-Cas in mobile genetic elements: counter-defence and beyond. Nat Rev Microbiol. 17, 513–525

30 Nishimura, T., Fakim, H., Brandmann, T., Youn, J. Y., Gingras, A. C., Jinek, M. and Fabian, M. R. (2018) Human MARF1 is an endoribonuclease that interacts with the DCP1:2 decapping complex and degrades target mRNAs. Nucleic Acids Res. 46, 12008–12021

31 Jumper, J., Evans, R., Pritzel, A., Green, T., Figurnov, M., Ronneberger, O., Tunyasuvunakool, K., Bates, R., Zidek, A., Potapenko, A., Bridgland, A., Meyer, C., Kohl, S. A. A., Ballard, A. J., Cowie, A., Romera-Paredes, B., Nikolov, S., Jain, R., Adler, J., Back, T., Petersen, S., Reiman, D., Clancy, E., Zielinski, M., Steinegger, M., Pacholska, M., Berghammer, T., Bodenstein, S., Silver, D., Vinyals, O., Senior, A. W., Kavukcuoglu, K., Kohli, P. and Hassabis, D. (2021) Highly accurate protein structure prediction with AlphaFold. Nature. 596, 583–589

32 Mirdita, M., Schutze, K., Moriwaki, Y., Heo, L., Ovchinnikov, S. and Steinegger, M. (2022) ColabFold: making protein folding accessible to all. Nat Methods. 19, 679–682

33 Holm, L., Laiho, A., Toronen, P. and Salgado, M. (2023) DALI shines a light on remote homologs: One hundred discoveries. Protein Sci. 32, e4519

34 Carte, J., Pfister, N. T., Compton, M. M., Terns, R. M. and Terns, M. P. (2010) Binding and cleavage of CRISPR RNA by Cas6. RNA. 16, 2181–2188

35 Hatoum-Aslan, A., Maniv, I. and Marraffini, L. A. (2011) Mature clustered, regularly interspaced, short palindromic repeats RNA (crRNA) length is measured by a ruler mechanism anchored at the precursor processing site. Proc Natl Acad Sci U S A. 108, 21218–21222

36 Hatoum-Aslan, A., Maniv, I., Samai, P. and Marraffini, L. A. (2014) Genetic characterization of antiplasmid immunity through a type III-A CRISPR-Cas system. J Bacteriol. 196, 310–317

37 Sokolowski, R. D., Graham, S. and White, M. F. (2014) Cas6 specificity and CRISPR RNA loading in a complex CRISPR-Cas system. Nucleic Acids Res. 42, 6532–6541

38 Zhang, J., Rouillon, C., Kerou, M., Reeks, J., Brugger, K., Graham, S., Reimann, J., Cannone, G., Liu, H., Albers, S. V., Naismith, J. H., Spagnolo, L. and White, M. F. (2012) Structure and mechanism of the CMR complex for CRISPR-mediated antiviral immunity. Mol Cell. 45, 303–313

39 Osawa, T., Inanaga, H., Sato, C. and Numata, T. (2015) Crystal structure of the CRISPR-Cas RNA silencing Cmr complex bound to a target analog. Mol Cell. 58, 418–430

40 Walker, F. C., Chou-Zheng, L., Dunkle, J. A. and Hatoum-Aslan, A. (2017) Molecular determinants for CRISPR RNA maturation in the Cas10-Csm complex and roles for non-Cas nucleases. Nucleic Acids Res. 45, 2112–2123

41 Chou-Zheng, L. and Hatoum-Aslan, A. (2022) Critical roles for ‘housekeeping’ nucleases in type III CRISPR-Cas immunity. Elife. 11 e81897.

42 Chou-Zheng, L. and Hatoum-Aslan, A. (2019) A type III-A CRISPR-Cas system employs degradosome nucleases to ensure robust immunity. Elife. 8 e45393.

43 Hale, C. R., Zhao, P., Olson, S., Duff, M. O., Graveley, B. R., Wells, L., Terns, R. M. and Terns, M. P. (2009) RNA-guided RNA cleavage by a CRISPR RNA-Cas protein complex. Cell. 139, 945–956

44 Tamulaitis, G., Kazlauskiene, M., Manakova, E., Venclovas, C., Nwokeoji, A. O., Dickman, M. J., Horvath, P. and Siksnys, V. (2014) Programmable RNA shredding by the type III-A CRISPR-Cas system of Streptococcus thermophilus. Mol Cell. 56, 506–517

45 Wang, J. Y., Hoel, C. M., Al-Shayeb, B., Banfield, J. F., Brohawn, S. G. and Doudna, J. A. (2021) Structural coordination between active sites of a CRISPR reverse transcriptase-integrase complex. Nat Commun. 12, 2571

46 Rouillon, C., Athukoralage, J. S., Graham, S., Gruschow, S. and White, M. F. (2019) Investigation of the cyclic oligoadenylate signaling pathway of type III CRISPR systems. In Crispr-Cas Enzymes (Bailey, S., ed.). pp. 191–218

47 Brown, T. S. and Bevilacqua, P. C. (2005) Method for assigning double-stranded RNA structures. Biotechniques. 38, 368, 370, 372

48 Bydovskii, E. I. and Klebanova, L. M. (1967) [Alkaline hydrolysis of ribonucleic acids]. Vopr Med Khim. 13, 299–303

49 Yang, W. (2011) Nucleases: diversity of structure, function and mechanism. Q Rev Biophys. 44, 1–93

